# MTAP deficiency is a novel biomarker in neuroendocrine neoplasms of the lung

**DOI:** 10.64898/2026.06.02.729487

**Authors:** Magdalena M. Brune, Luca Roma, Nikolaus Deigendesch, Felix Meissner, Sarp Uzun, Jürgen Hench, Obinna Chijioke, Ivana Bratic Hench, Jumpei Kashima, Petra Hirschmann, Julian Pollinger, Keith M. Kerr, David König, Chantal Pauli, Sebastian R. Ott, Spasenija Savic Prince, Martina Haberecker, Lukas Bubendorf

**Author notes:** These authors contributed equally to this work (Magdalena M. Brune, Luca Roma). Corresponding author: Address for correspondence: Lukas Bubendorf, MD, PhD. Institute of Medical Genetics and Pathology, University Hospital Basel, Schönbeinstrasse 40, 4031 Basel, Switzerland. No funding was received for conducting this study.

## Abstract

**Introduction:** MTAP emerges as potential predictive biomarker for MTA-cooperative PRMT5-inhibitors. Although MTAP attracts increasing attention in non-small cell lung cancer, its role in pulmonary neuroendocrine neoplasms (NENs) remains largely unexplored.

**Methods:** Here, we assessed the prevalence of MTAP deficiency in 209 pulmonary NENs using immunohistochemistry (IHC). Additionally, we performed fluorescence in situ hybridization (FISH), whole exome sequencing (WES), deep proteomic profiling, transcriptomic, and methylation analyses of selected MTAP deficient and proficient carcinoids to further elucidate the underlying mechanisms of MTAP expression pattern.

**Results:** MTAP deficiency by IHC was detected in all neuroendocrine precursor lesions (n=17), 92% of typical carcinoids (n=51), 86% of atypical carcinoids (n=21), and 10% of large cell neuroendocrine carcinomas (LCNEC) (n=30). In contrast, all small cell lung cancers (SCLC) were MTAP proficient (n=90). In MTAP deficient carcinoids, FISH and WES did not detect homozygous 9p21 deletions, and methylation analysis showed no evidence of *MTAP* promoter hypermethylation.

Comparing MTAP deficient and MTAP proficient carcinoids, proteomic data showed a clear separation between the two groups. Further, there was an inverse correlation between the expression of MTAP and OTP (orthopedia homeobox protein), which is known as a strong prognostic marker in pulmonary carcinoids.

**Discussion:** MTAP deficiency is a novel hallmark of neuroendocrine precursors and most pulmonary carcinoids, clearly distinguishing the latter from SCLC and most LCNEC. It is neither caused by 9p21 deletion, nor by MTAP promoter hypermethylation. MTAP deficiency is a group-defining feature of carcinoids that might pave the way for new therapeutic approaches.

## 1. Introduction

Deciphering lung cancer heterogeneity and identifying therapeutic targets remains a major challenge in thoracic oncology. However, advances in sequencing technology over the last two decades have significantly improved our understanding of tumor biology, paving the way for the development of novel and more effective treatment strategies^1^. Targeted therapies against oncogenic drivers and immune checkpoint inhibitors (ICI) significantly improved the management of non-small cell lung cancer (NSCLC). In parallel, therapeutic options for neuroendocrine neoplasms (NENs) of the lung evolved significantly during the last decade^2^. NENs comprise approximately 20% of all pulmonary malignancies and encompass a variety of different entities, ranging from well-differentiated typical carcinoids to highly aggressive small cell lung cancers (SCLC) and large cell neuroendocrine carcinomas (LCNEC)^3^. Carcinoids of the lung account for 1-2% of all lung tumors and occur slightly more often in women^4^. In lung carcinoids, surgery is the treatment of choice in localized disease in both typical and atypical carcinoids^4^. Nevertheless, recurrences are common, and treatment of systemic disease includes endocrine, cytotoxic and radiotherapeutic strategies^5^. Moreover, 3-year overall survival for patients with metastatic atypical carcinoids is only about 22%, and treatment efficacy varies considerably^6^. Currently, targeting S-methyl-5’-thioadenosine phosphorylase (MTAP) deficient tumors has emerged as a promising novel approach in NSCLC^7^. Loss of MTAP expression has been described across various tumor types^7,8,9^ and is associated with vulnerability to synthetic lethality-based approaches targeting PRMT5 or MAT2A^7^. Recent studies suggest that loss of MTAP expression leads to intracellular accumulation of its substrate, methylthioadenosine (MTA), which partially suppresses PRMT5 activity. This suppression impairs PRMT5 function and ultimately induces cell cycle arrest and apoptosis^10^. As a result, MTAP deficient cells become highly reliant on the residual activity of PRMT5 for survival. Notably, further accumulation of MTA selectively compromises cell viability in MTAP deficient cells^10^, rendering them vulnerable to synthetic lethality–based therapeutic strategies^11^. MTAP deficiency is commonly caused by homozygous deletions of the 9p21 chromosomal region, one of the most frequent copy number variations (CNV) in human cancer^12,13,14^, and it often co-occurs with loss of the directly adjacent *CDKN2A/B* (cyclin-dependent kinase inhibitor 2A/B) locus. We and others have recently studied MTAP deficiency in NSCLC^15,16^, but its prevalence and diagnostic utility in other neoplasms such as pulmonary NENs remain largely unexplored^8^. Given the increasing interest in MTAP as a prognostic and predictive biomarker, we systematically evaluated its protein expression by immunohistochemistry (IHC) across a comprehensive cohort of lung NENs and neuroendocrine precursor lesions, highlighting a novel diagnostic and potentially predictive role for MTAP loss in this tumor spectrum. In addition, we integrated whole exome sequencing (WES), bulk RNAseq, methylation and proteomic data to further explore *MTAP* expression patterns.

## 2. Results

### 2.1 MTAP deficiency is present in all neuroendocrine precursor lesions and most carcinoids

We found MTAP deficiency by IHC (Figure 1A-C, Table 1) in all neuroendocrine precursor lesions (n=17), in 47/51 (92%) of typical carcinoids and in 18/21 (86%) of atypical carcinoids. In contrast, none of the SCLC (n=90) and only 3/30 (10%) of the LCNEC were MTAP deficient (Figure 1E, F).

**Figure 1:**
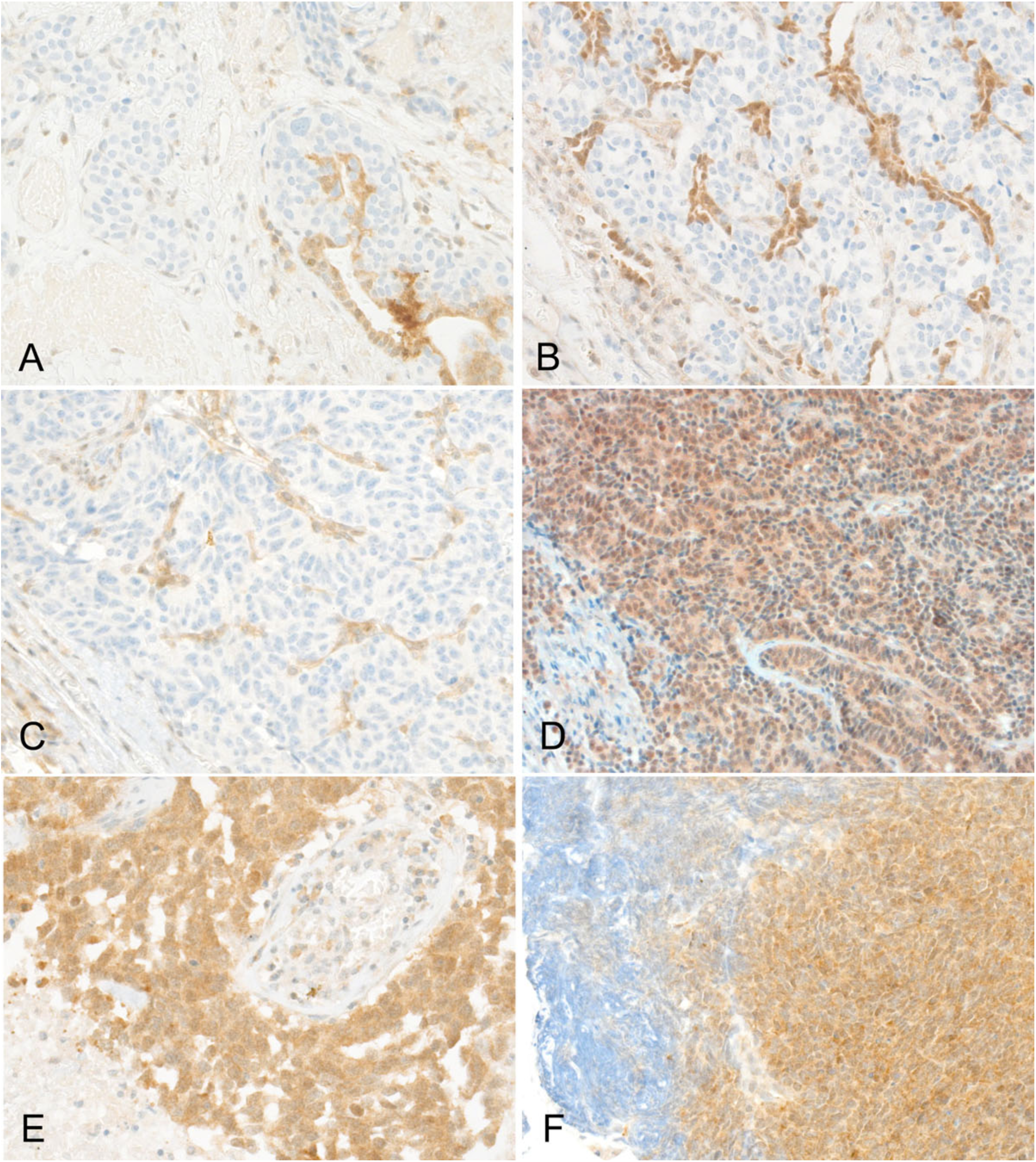
MTAP expression status in neuroendocrine lesions and neoplasia by immunohistochemistry. A) MTAP deficient tumorlet (x400), B) MTAP deficient typical carcinoid with residual MTAP proficient benign type 2 alveolar cells (x400), C) MTAP deficient atypical carcinoid with numerous capillaries serving as internal positive staining control (x400), C) Typical carcinoid D) MTAP proficient typical carcinoid (x200) E) MTAP proficient large cell neuroendocrine carcinoma (x400), F) MTAP proficient small cell carcinoma. Note the impaired staining of area with crush artifact (x400).

**Table 1.** Clinical parameters, specimen types, tumor and clinical stage, smoking history and MTAP expression by immunohistochemistry cM0, localized disease; cM1, distant metastases present; IHC, immunohistochemistry; LCNEC, large cell neuroendocrine carcinoma; MTAP, S-methyl-5’-thioadenosine phosphorylase; NA, not applicable; NE, neuroendocrine; pT, pathologic T-classification; SCLC, small-cell lung cancer.

| *Patient Cohort* | **All cases**<br>(n=209) (%) | **NE precursors**<br>(n=17) (%) | **Typical carcinoids**<br>(n=51) (%) | **Atypical carcinoids**<br>(n=21) (%) | **SCLC**<br>(n=90) (%) | **LCNEC**<br>(n=30) (%) |
| --- | --- | --- | --- | --- | --- | --- |
| **Age, median (IQR)** | 67 (59-74) | 68 (59-74) | 63 (48-73) | 66 (55-72) | 67.5 (61-73.75) | 66.5 (60.5-74) |
| **Sex** |  |  |  |  |  |  |
| Male | 81 (38.6) | 3 (17.6) | 19 (37.3) | 10 (47.6) | 36 (40.0) | 13 (43.3) |
| Female | 128 (61.4) | 14 (82.4) | 32 (62.7) | 11 (52.4) | 54 (60.0) | 17 (56.6) |
| **Specimen type** |  |  |  |  |  |  |
| Biopsy | 80 (38.3) | 0 (0.0) | 0 (0.0) | 0 (0.0) | 79 (87.8) | 1 (3.3) |
| Resection | 129 (61.7) | 17 (100.0) | 51 (100.0) | 21 (100.0) | 11 (12.2) | 29 (96.7) |
| **T-classification** |  |  |  |  |  |  |
| pT1 | NA | NA | 41 (80.4) | 12 (57.1) | NA | 12 (40.0) |
| pT2 | NA | NA | 10 (19.6) | 6 (28.6) | NA | 11 (36.7) |
| pT3 | NA | NA | 0 (0.0) | 2 (9.5) | NA | 3 (10.0) |
| pT4 | NA | NA | 0 (0.0) | 1 (4.8) | NA | 2 (6.7) |
| unknown | NA | NA | 0 (0.0) | 0 (0.0) | NA | 2 (6.7) |
| **M-classification** |  |  |  |  |  |  |
| cM0 | NA | NA | 50 (98.0) | 20 (95.2) | NA | 25 (83.3) |
| cM1 | NA | NA | 0 (0.0) | 1 (4.8) | NA | 2 (6.7) |
| unknown | NA | NA | 1 (2.0) | 0 (0.0) | NA | 3 (10.0) |
| **Smoking** |  |  |  |  |  |  |
| never smoker | NA | NA | 22 (43.1) | 9 (42.6) | NA | 0 (0.0) |
| former smoker | NA | NA | 23 (45.1) | 7 (33.3) | NA | 14 (46.7) |
| current smoker | NA | NA | 5 (9.8) | 4 (19.0) | NA | 10 (33.3) |
| unknown | NA | NA | 1 (2.0) | 1 (4.8) | NA | 6 (20.0) |
| **MTAP by IHC** |  |  |  |  |  |  |
| proficient | NA | 0 (0.0) | 4 (7.8) | 3 (14.3) | 90 (100.0) | 27 (90.0) |
| deficient | NA | 17 (0.0) | 47 (92.2) | 18 (85.7) | 0 (0.0) | 3 (10.0) |

In our cohort of lung carcinoids (n=72) (Table 1), MTAP deficiency was more frequent in females than in males (Fisher’s exact test, *p* < 0.01). Beyond that, there was no significant association between MTAP expression and age, smoking history, clinical stage, tumor size, pathological T classification or the expression of TTF-1, chromogranin, or synaptophysin by IHC. Data on TTF-1 were available in 57% (41/72), on synaptophysin (SYP) in 92% (66/72), and on chromogranin A (CHGA) in 67% (48/72).

To further elucidate the underlying mechanism for MTAP deficiency in carcinoids, twelve MTAP deficient carcinoids were analyzed by FISH, but none of these revealed a homozygous deletion of the chromosomal region 9p21 (Figure 2). MTAP proficiency by IHC in SCLC and LCNEC was confirmed by *MTAP* RNA expression analysis in two publicly available datasets (Supplementary Figure 1A, Supplementary Figure 1B, Material and Methods), which both showed *MTAP* expression levels similar to those of the neuroendocrine markers *CHGA, SYP* and *NCAM1/CD56*. In contrast, the evaluation of a previously published dataset of 30 lung carcinoids (containing 17 typical and 13 atypical carcinoids)^17^ showed that *MTAP* RNA expression levels in carcinoids were significantly lower compared to the expression of established neuroendocrine markers, including chromogranin A, synaptophysin, and *NCAM1*/*CD56* (*p* < 0.01) (Supplementary Figure 1A).

**Figure 2:**
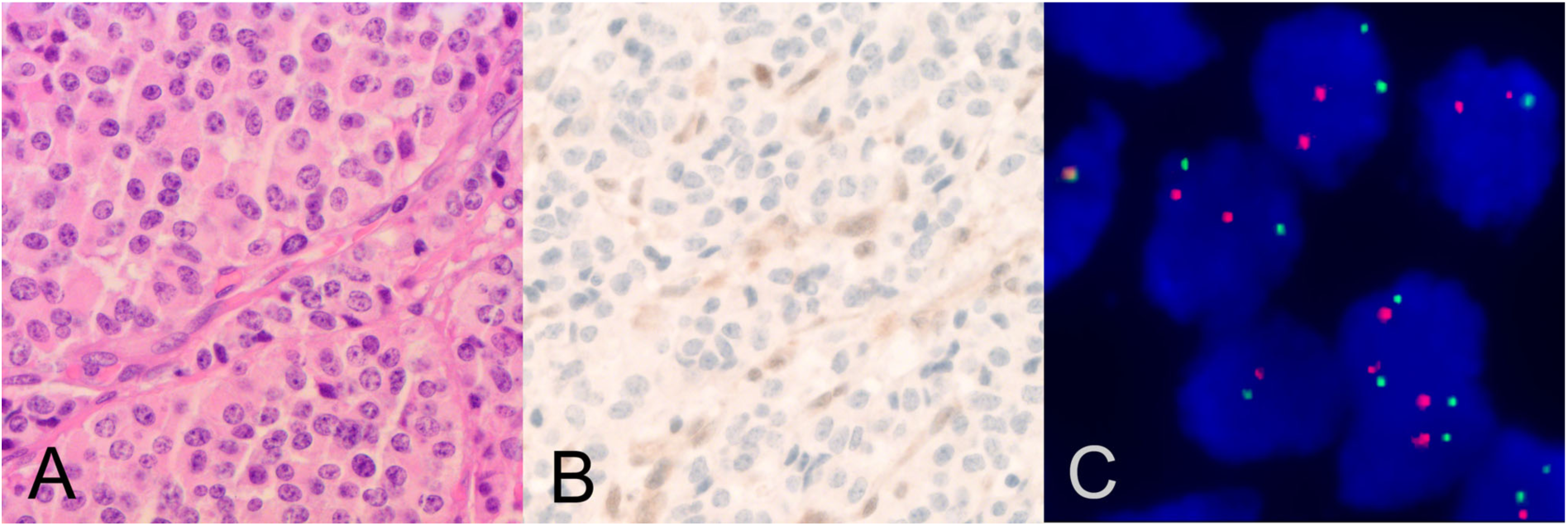
MTAP expression status and fluorescence in situ hybridization. A)-C) MTAP deficient (typical) carcinoid intact without CDKN2A/MTAP deletion A) Hematoxylin-eosin staining (H&E, x400), B) MTAP by IHC (x400), C) Tumor cell showing red centromeric 9 signals intact green 9p21/CDKN2A/MTAP signals (x1000).

### 2.2 Whole exome sequencing detects no copy number variations at 9p21

Macrodissected tumor areas of 6 MTAP proficient and 5 MTAP deficient carcinoids and matched non-tumor samples were subjected to WES (Supplementary Figure 2A, Supplementary Table 1-4). No copy number variations were detected at chromosome 9p21.3, which harbors *MTAP* and the adjacent *CDKN2A/B,* confirming our FISH results (Supplementary Figure 2B), neither in MTAP deficient nor in MTAP proficient tumors. The average tumor mutational burden (TMB) was 3 mutations/Mb (range 0.6 – 8), with MTAP proficient samples showing a higher TMB than MTAP deficient (4.1 vs 1.6 mutation/Mb respectively) (Supplementary Table 3). At the gene level, *ROS1*, *NLGN3* and *RYR2* were the most frequently altered genes (2/11 patients, 18%). *TP53* and *MEN1* mutations were detected only in two distinct MTAP proficient patients. Functional pathway grouping highlighted targeted alterations in specific key cellular processes: we detected a clonal nonsense mutation in the SWI/SNF complex subunit *ARID1A* (p.Gln1499*), and the MAPK/ERK signaling cascade harbored variants in established oncogenic drivers, including *BRAF* (p.Gly509Ser), *HRAS* (p.Gln61Arg) and *MET* p.Ala1372_Asn1376del (Supplementary Table 2 and Supplementary Figure 2A). In general, we found a high degree of inter-patient genomic heterogeneity and could not find any difference between MTAP proficient and MTAP deficient samples regarding the presence and kind of mutations.

### 2.3 Methylation analysis shows no MTAP promoter hypermethylation

To investigate whether the observed loss of MTAP expression was driven by epigenetic alterations, we performed DNA methylation profiling. Analyses were conducted on five carcinoid tumor samples (two MTAP proficient, three MTAP deficient) with available fresh-frozen material, integrated with matching WES, transcriptomic, and proteomic data. We specifically examined the methylation status of CpG sites within the promoter regions of *MTAP* and *OTP (*orthopedia homeobox protein). Regarding *MTAP*, the promoter region appeared generally hypomethylated across the entire cohort (Supplementary Figure 2D). Specifically, the analysis of the *MTAP* promoter-associated probe cg25162921 revealed no significant differential methylation between MTAP proficient and MTAP deficient samples (Student’s t-test, *p* = 0.6), suggesting that DNA promoter hypermethylation is unlikely to be the primary mechanism underlying MTAP loss in these tumors (Supplementary Figure 2E). This finding was further validated by analyzing two independent lung neuroendocrine tumor methylation cohorts^17,18^ (*Alcala et al*. and *Laddha et al*.), which confirmed that none of the carcinoid samples exhibited promoter hypermethylation of MTAP (Supplementary Figures 2F and 2G). Conversely, we identified a significant epigenetic divergence in the *OTP* promoter region. The CpG probe cg02493167, located close to the *OTP* promoter, showed a significant difference in methylation levels between the two groups (Student’s t-test, *p* = 0.027), with hypomethylation observed in MTAP-deficient samples (Supplementary Figure 2E).

### 2.4 Proteomic profiling reveals distinct molecular features associated with MTAP status

Deep proteomic profiling following laser microdissection was performed on eight resected FFPE carcinoid tumor samples, comprising four MTAP deficient and four MTAP proficient cases. Tumor regions were annotated on H&E-stained sections, with cumulative areas of approximately 30,000 µm², corresponding to approximately 300 to 500 tumor cells. Regions were carefully delineated to minimize co-sampling of stromal components and infiltrating immune cells, yielding near tumor-pure proteomic profiles^19^. The spectral library comprised 167,826 precursors and 9,223 protein groups; after filtering, 7,526 proteins were consistently quantified across the dataset.

Unsupervised principal component analysis (PCA) demonstrated a clear separation between MTAP deficient and MTAP proficient tumors, consistent with transcriptomic patterns observed in the subset of five specimens with matched RNA-seq data (see below, Figure 3A). Notably, MTAP deficient tumors exhibited lower inter-sample heterogeneity at the proteomic level compared to MTAP proficient tumors, suggesting that MTAP deficient carcinoids represent a more molecularly homogeneous subgroup.

**Figure 3:**
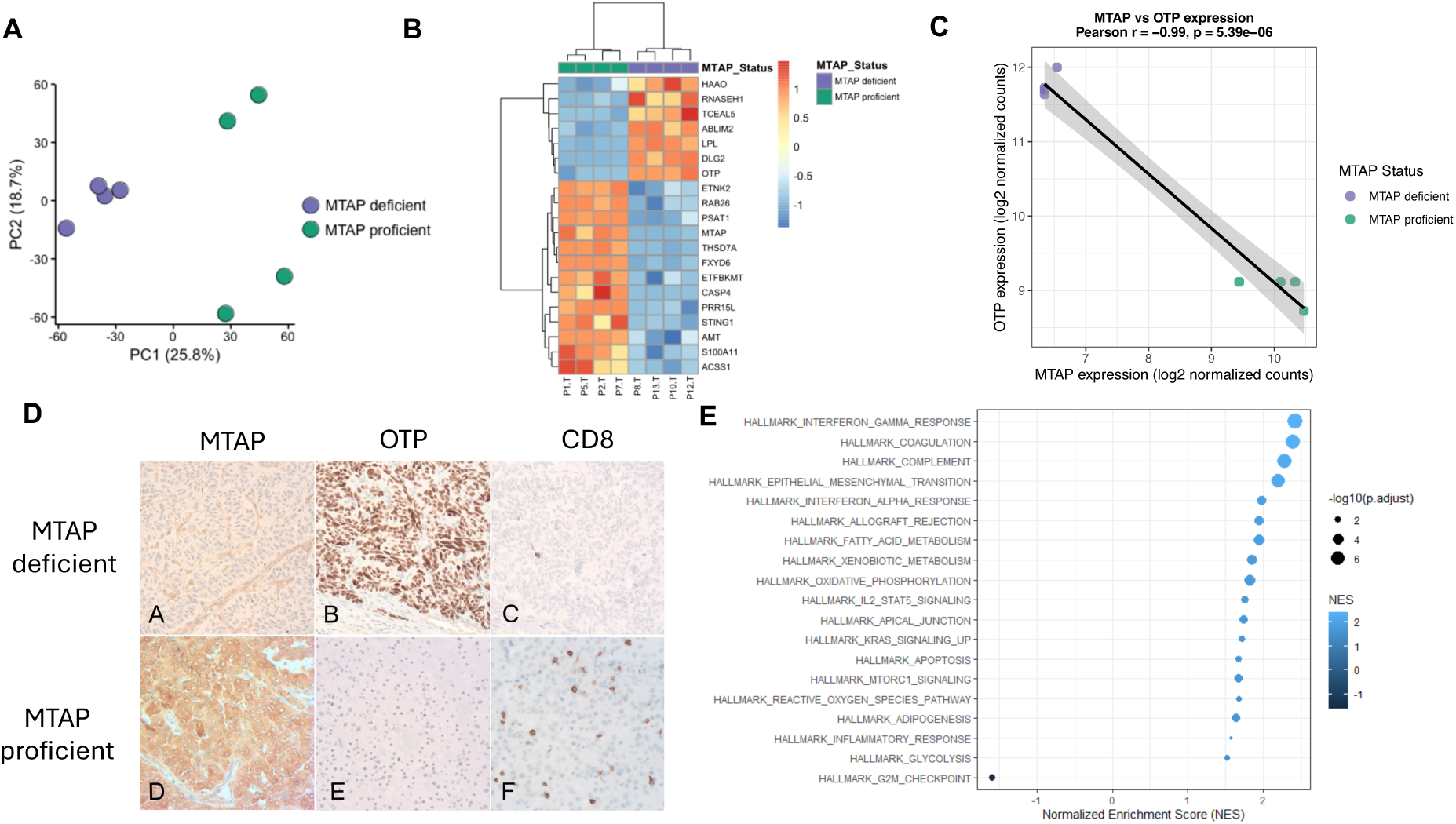
Proteomic, and immunohistochemical characterization of MTAP proficient versus MTAP deficient carcinoids. **A)** Principal component analysis (PCA) showing proteomics profiles based on all genes. **B)** Unsupervised heatmap showing the differentially expressed proteins in carcinoids comparing MTAP proficient (n=4, right) versus MTAP deficient samples (n=4, left) by proteomics. **C)** Scatterplot showing *Pearson’s* correlation between *MTAP* and *OTP* expression in proteomics comparing MTAP proficient (n=4) versus MTAP deficient samples (n=4). *Pearson’s* correlation coefficient *(r*) and *p* are indicated. **D)** Correlation of MTAP status with OTP status and immune-infiltration by CD8+ T-cells (x200, each). A) MTAP deficient typical carcinoid with B) OTP expression and C) cold tumor microenvironment with virtual lack of infiltrating CD8+ T-cells. D) MTAP proficient atypical carcinoid E) without OTP expression D) and immune-infiltration by CD8+ T-cells. **E)** Gene set enrichment analysis (GSEA) showing hallmark pathways in MTAP proficient versus MTAP deficient samples based on proteomics. Significantly deregulated pathways (adjusted *p <* 0.05) are highlighted blue, and the size of the dots is proportional to the significance of the statistical test.

Differential protein expression analysis using the Limma package identified 71 proteins significantly upregulated, and 30 proteins significantly downregulated in MTAP proficient tumors relative to MTAP deficient counterparts (log2 Fold Change > 1, FDR-adjusted *p* < 0.05) (Supplementary Figure 3A). Among the upregulated proteins, S100A11 showed the strongest increase. MTAP was additionally found to be significantly upregulated. In contrast, downregulated proteins included LPL, DLG2 and OTP (log2 Fold Change < –1, FDR-adjusted *p* < 0.05) (Supplementary Table 6; Figure 3B and Supplementary Figure 3A). Missing values were imputed using local stochastic imputation around detected values to account for proteins detected only in a limited number of samples or completely absent in either the MTAP-proficient or –deficient group. Consequently, the calculated fold changes and adjusted *p* values of these proteins are strongly influenced by the imputation strategy and should be interpreted qualitatively rather than quantitatively in this cohort with limited sample size.

### 2.5 Transcriptomics findings confirm distinct molecular features associated with MTAP status

To investigate transcriptional alterations associated with MTAP status, bulk RNA sequencing (RNA-seq) was performed on five carcinoid tumor samples matching to WES and proteomics data (n=2 MTAP proficient, n=3 MTAP deficient), for which fresh-frozen tumor material was available. Unsupervised principal component analysis (PCA) based on the 100 most variable genes showed clear separation of MTAP deficient and MTAP proficient samples, suggesting two distinct transcriptomic profiles, consistent with proteomic patterns (Supplementary Figure 3B). Differential expression analysis comparing the two groups identified 968 upregulated and 736 downregulated genes in MTAP proficient samples compared to MTAP deficient tumors. Among the genes upregulated in MTAP proficient samples were *MTAP* (log2 Fold Change = 3.2 and adjusted *p* < 0.01), *CDKN2A* (log2 Fold Change = 4.6 and adjusted *p* < 0.01) and *S100A11* (log 2 Fold Change = 3.1 and adjusted *p* = 0.02), whereas *OTP* was among the downregulated genes (log2 Fold Change = –16 and adjusted *p* < 0.01) (Supplementary Table 5 and Supplementary Figure 3C).

### 2.6 MTAP expression is significantly associated with OTP expression

Proteomic analysis revealed a strong inverse correlation between MTAP and OTP expression (*Pearson’s* r = −0.99, *p* < 0.01; Figure 3C). This relationship was also found at the transcriptomic level (r = −0.93, *p* = 0.02; Supplementary Figure 3D) and was further supported by an independent publicly available transcriptomic dataset of pulmonary carcinoids^20^, which showed a significant negative correlation between *MTAP* and *OTP* expression (r = −0.52, *p* < 0.01; Supplementary Figure 3D). The downregulation of OTP depending on MTAP status was further confirmed by IHC, as all investigated MTAP proficient samples (n = 6) were completely negative by IHC for OTP, whereas all tested MTAP deficient samples (n = 8) showed consistent OTP expression (Figure 3D).

### 2.7 Pathway analysis reveals enrichment of immune-related pathways in MTAP proficient carcinoids

Pathway analysis of dysregulated genes derived from proteomic and transcriptomic data revealed enrichment of immune-related pathways, including HALLMARK_INTERFERON_GAMMA_RESPONSE and HALLMARK_INTERFERON_ALPHA_RESPONSE in MTAP proficient samples (Figure 3E and Supplementary Figure 3E), suggesting increased immune-associated transcriptional activity in this subgroup.

These findings were supported by the deconvolution method applied to estimate immune cell infiltration from bulk RNA-seq data, which consistently yielded higher immune infiltration scores in MTAP proficient samples (Supplementary Figure 3F). To further validate these observations, CD8 immunohistochemistry was performed in MTAP proficient (n=6) and MTAP deficient (n=7) samples. The percentage of CD8 positive immune cells tended to be higher in MTAP proficient samples compared to MTAP deficient samples, although this difference was not statistically significant (Figure 3D and Supplementary Figure 3G).

### 2.8 Carcinoids with heterogeneous MTAP expression

In four additional carcinoids (two typical and two atypical), we observed heterogeneous MTAP expression (Figure 4A). These cases were excluded from the statistical analysis but are described here and in Figure 4. All cases were pT1 lung resection specimens from male patients and were classified as MTAP deficient, with loss of MTAP expression in at least 50% of tumor cells. One specimen was suitable for further comparative IHC analysis because it contained well-demarcated MTAP proficient and MTAP deficient areas (Figure 4A, B). Infiltration by CD8 positive immune cells was clearly higher in the MTAP proficient than in the MTAP deficient areas (Figure 4C), recapitulating the association between MTAP expression and immune infiltration observed at the cohort level.

**Figure 4:**
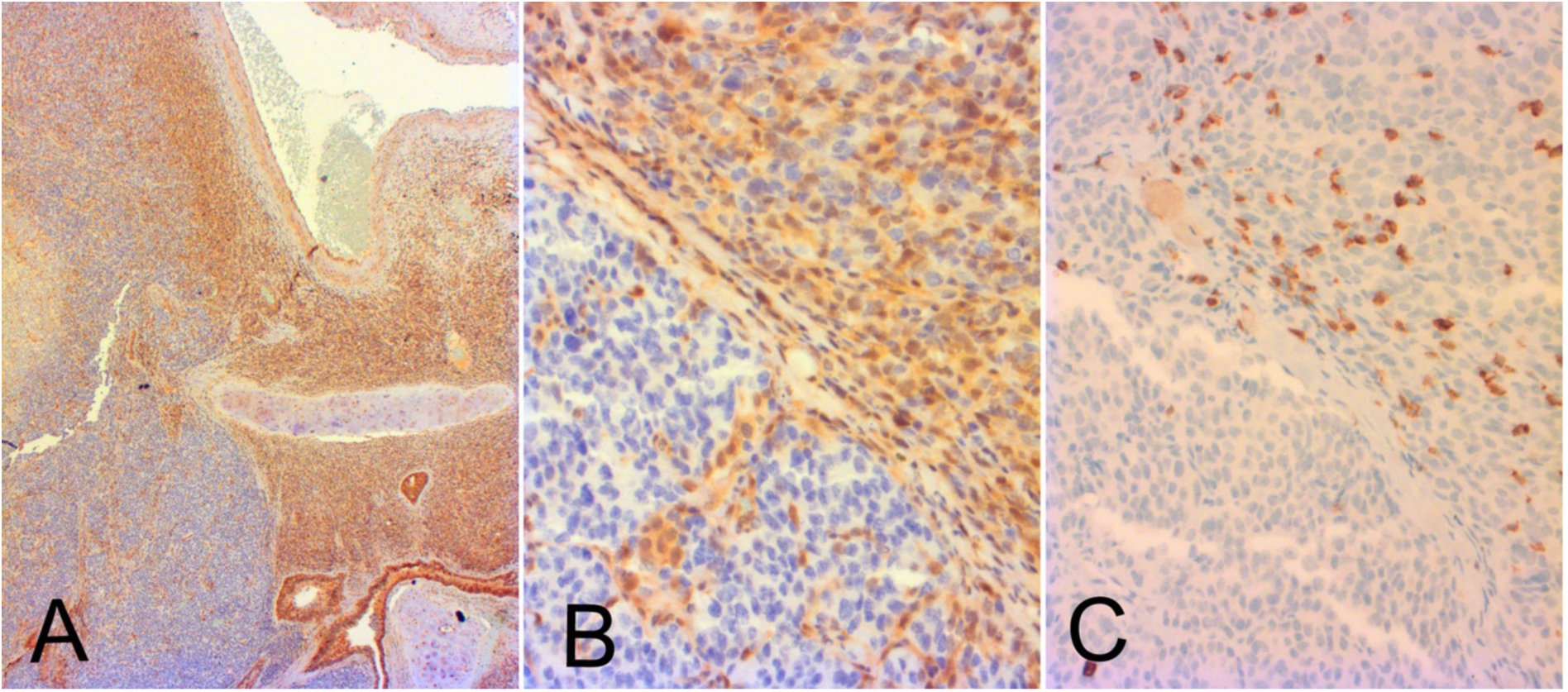
Atypical carcinoid with heterogenous MTAP expression and CD8+ T-cell infiltration. **A)** Clearly defined tumor areas with MTAP deficiency (upper half) and MTAP proficiency (lower half) including bronchial mucosa with cartilage and blood vessel (overview, x25). **B)** Sharp border between MTAP deficient (upper part) and lower MTAP proficient (lower part) tumor area (x400). **C)** high immune cell infiltration restricted to MTAP proficient tumor area (x400). A-B): MTAP MSVA 741R antibody on Leica BOND-PRIME automated immunostainer was used for this IHC.

## 3. Discussion

MTAP deficiency associated with homozygous 9p21 deletion is currently receiving considerable attention as a predictive biomarker in NSCLC and other solid tumors^15,16,21,22^. Here, we show that MTAP deficiency by IHC is a novel hallmark of pulmonary carcinoids and their precursor lesions, with similarly high prevalence in typical and atypical carcinoids (86-92%), yet without underlying 9p21 deletion.

The high prevalence of MTAP deficiency in pulmonary carcinoids has remained under the radar of previous molecular profiling studies of lung NENs^17,18,20,23,24,25,26,27,28,29^. This can mainly be attributed to the absence of homozygous 9p21 deletion in lung carcinoids, which is commonly used as a surrogate for MTAP loss in other tumor types. Furthermore, uniform MTAP expression in benign tissues can mask MTAP loss in expression profiling studies, especially in specimens with relatively low tumor content. The importance of protein-level assessment is underscored by recent tissue microarray studies demonstrating frequent MTAP loss in pulmonary neuroendocrine tumors despite the absence of 9p21 deletion^8^. The mechanism of MTAP silencing in carcinoids remains unclear. Since we found no evidence for promoter hypermethylation in two independent datasets of lung carcinoids^17,18^ (A*lcala et al.,* 55 samples; *Laddha et al.,* 18 samples) and our own analyses, promoter methylation is unlikely to regulate MTAP expression. Unfortunately, data on MTAP expression by IHC is not available for the above-mentioned datasets. Nevertheless, the limited probe coverage of Infinium methylation arrays within gene regulatory regions represents an important technical limitation, as biologically relevant CpG sites may not be sufficiently captured and therefore subtle regulatory methylation changes cannot be entirely excluded. It is possible that not yet identified epigenetic or post-transcriptional regulatory mechanisms, such as DNA demethylation, chromatin remodeling, long non-coding RNAs (lncRNAs) or microRNAs (miRNAs) may influence MTAP expression. This would also explain the lack of a significant difference between MTAP proficient and MTAP deficient samples when comparing the mutational landscape (Supplementary Figure 2A).

Interestingly, comparative proteomic and transcriptomic analyses, together with IHC, revealed an almost perfect but inverse correlation between MTAP and OTP expression, evoking a “chicken or egg” causality dilemma. Other than MTAP deficiency, OTP expression is long known and unique to pulmonary carcinoids and not expressed in other neuroendocrine tumors^30,31^. However, there is currently no published evidence of a direct functional connection between these two proteins. One could speculate that the reduced PRMT5 in MTAP deficient pulmonary carcinoids might lead to an upregulation of *OTP* gene expression through altered histone methylation^32^. Interestingly, our data and that of others show that OTP high (corresponding to MTAP deficient) pulmonary carcinoids harbor a lower methylation level than OTP low (corresponding to MTAP proficient) carcinoids, suggesting that MTAP deficiency might alter DNA methylation by decreased PRMT5 activity^30^. Additionally, transcriptomic and proteomic analyses identified S100A11 as one of the upregulated proteins in MTAP proficient tumors, again with no published data on the possible functional connection between these two proteins.

Our findings have diagnostic, prognostic and potential therapeutic implications in patients with neuroendocrine neoplasms of the lung. Since SCLC was consistently MTAP proficient in our study, detection of MTAP deficiency could be used to differentiate carcinoids from SCLC in problematic small-sized diagnostic samples, thus complementing Ki67 analysis that is currently often used for this purpose^33^. Homozygous MTAP deletion is associated with poor prognosis across different solid tumor types, with a large real-world study showing significantly reduced overall survival in patients with MTAP deleted solid tumors compared to those with MTAP proficient tumors^34^. In lung carcinoids, our data indirectly point to the opposite direction. The role of OTP has extensively been analyzed by several research groups showing an adverse prognosis of the small subgroup of OTP-low carcinoids^26,31,35,36^. This is in line with the generally good prognosis of most pulmonary carcinoids with only a relatively small proportion showing adverse outcomes^31,37^.

Physiologically, the nuclear transcription factor *OTP* is involved in hypothalamic development and differentiation^38^ and it has gained attention as a diagnostic feature of pulmonary carcinoids with nuclear expression in 80% of the cases and a strong prognostic impact^39^. Lately, OTP expression has been used to identify the recently proposed molecular subgroups A1 and A2 of pulmonary carcinoids that displayed a lower risk of recurrence and higher expression of key therapeutic targets compared to the OTP-low subgroup B^18,27^. OTP’s inverse correlation with MTAP infers an equally adverse prognostic effect of MTAP proficiency in lung carcinoids, as these cases likely correspond to subgroup B^26,33,36^. Therefore, MTAP proficiency in pulmonary carcinoids potentially defines a subgroup of 10-20% of lung carcinoids that might require particular attention beyond the current classification into typical and atypical carcinoids.

Pulmonary carcinoids with aggressive behavior are relatively rare but still present a relevant clinical challenge^40,41^. Given the limited therapeutic options with only moderate response rates, there is an apparent need for new therapeutic strategies^3,40,41^. Pre-clinical studies and clinical trials are needed to explore if MTA-cooperative PRMT5– or MAT2A-inhibitors could potentially represent a novel option for patients with unresectable or metastatic, MTAP deficient carcinoids. However, this might not apply to most aggressive carcinoids, as they are more likely to be MTAP proficient. Our study also supports the current notion that immunotherapy might serve as a future treatment option in a subgroup of lung carcinoids^40,41^. The combined pathway analysis of proteomic and transcriptomic data, together with IHC validation, consistently indicated enrichment of immune-related processes and increased immune cell infiltration in MTAP proficient tumors. This is in line with a link between MTAP status and the tumor immune microenvironment that is well-known from other solid malignancies^42,37^. Although the underlying biological mechanisms remain incompletely understood, accumulating MTA in the tumor seems to suppress CD8+ T-cell functions via the inhibition of PRMT5 and by adenosine receptor agonism which could at least partially explain the cold tumor microenvironment in MTAP deficient tumors^37^.

In our small series of 30 LCNEC, three were MTAP deficient, which is well in line with published data describing *CDKN2A* deletion in up to 8% of LCNEC^43,44,45^. All our SCLC were MTAP proficient by IHC, which is supported by RNA expression analysis of publicly available datasets (Supplementary Figure 1A)^46^. Most interestingly, *Rekhtman et al.* have recently proposed a new subset of atypical SCLC (aSCLC) that account for <5% of all SCLC, lack co-deletion of *TP53* and *RB1*, arise mostly in never/light smokers and seem to evolve from carcinoids^29^. It could be possible that such aSCLC might also be MTAP deficient without underlying 9p21.3 deletion akin to carcinoids. Indeed, publicly available data from this study at cBioPortal^47^ revealed lower expression of MTAP RNA in 2/7 aSCLC^29^. Additionally, rare focal deletions of 9p21.3 have been described in SCLC^48^. Potentially, these cases are also MTAP deficient at the protein level.

The proteomic datasets with matched transcriptomic data in a subset of cases represent, to our knowledge, one of the most comprehensive multi-omics characterizations of lung carcinoids available to date, with a very high tumor cell content and many identified proteins. Proteomic and transcriptomic analyses showed that MTAP deficient carcinoids form a distinct cluster, with less molecular heterogeneity compared to MTAP proficient tumors, identifiable across independent platforms. In addition, the integration of proteomic pathway analysis with transcriptomic data provides a robust and complementary view of the biological processes, such as immunoregulatory pathways, associated with MTAP status.

The low number of cases, due to the rarity of carcinoids and the very limited availability of fresh material suitable for bulk RNA sequencing and methylation microarray analyses is a limitation of our study, as more cases are needed to explore the association with other markers and recently proposed molecular carcinoid subtypes, and outcome^18,27,41^. Our illustrated carcinoid with heterogenous MTAP deficiency associated with different density of immune infiltration is impressive but anecdotal. Further studies are needed to define the prevalence, underlying biology and clinical significance of heterogeneous MTAP expression status in carcinoids.

In summary, MTAP deficiency is a novel hallmark of carcinoids and neuroendocrine precursor lesions and helps in the distinction towards SCLC and most LCNEC. MTAP deficiency in carcinoids is not caused by 9p21 deletion or *MTAP* promoter hypermethylation but is likely influenced by not yet fully elucidated epigenetic mechanisms. MTAP deficiency might pave the way for novel therapeutic approaches with MTA-cooperative PRMT5 and MAT2A inhibitors in MTAP deficient lung carcinoids, whereas patients with MTAP proficient carcinoids might be the main candidates for immunotherapy in future clinical trials.

## 4 Materials and Methods

### 4.1 Patient cohort

We retrospectively investigated MTAP expression by IHC in different NENs and neuroendocrine precursor lesions of the lung. Biopsy (38.3%) and resection (61.7%) specimens were retrieved from the archive of the Institute of Medical Genetics and Pathology, University Hospital Basel (USB), Switzerland, (from either internal routine diagnostics or referral cases sent to our institution), or from the Department of Pathology and Molecular Pathology, University Hospital Zurich (USZ), Switzerland or the Department of Pathology, Aberdeen University Medical School, University of Aberdeen (UA), United Kingdom. This study was approved by the local Ethics Committee (Ethikkomission Nordwestschweiz. Project-ID 2023-00843 and BASEC-2021-00417). The statistical analysis of the patient cohort was performed using Fisher’s exact test.

A total of 209 tissue samples were collected from 208 patients with NENs or precursor lesions of the lung, over a period from 2002 to 2026. The study included 90 samples of SCLC (all from USB), 30 samples of LCNEC (12 from USB and 18 from USZ), 72 samples of carcinoids (36 from USB and 36 from USZ, respectively, including 51 typical and 21 atypical), and 17 samples of neuroendocrine precursor lesions (10 from USB and 7 from UA, respectively), including seven tumorlets and eight diffuse idiopathic pulmonary neuroendocrine cell hyperplasia (DIPNECH) specimens, as well as two specimens with tumorlets and DIPNECH. A total of 188 samples were directly obtained from the lung, eleven from locoregional (thoracic) lymph node metastases, and nine from distant metastatic sites (liver, bone, brain, non-regional lymph nodes). The exact origin of one sample was not specified. The cohort included 128 female and 81 male patients. Their median age was 67 years (IQR 59-74 years).

Information about TNM-stage according to the Union of International Cancer Control (UICC, 8th Edition) was collected for patients with carcinoids and LCNEC samples. Data were available in 97.2% (70/72) of carcinoid and in 90.0% (27/30) of LCNEC patients. Additionally, smoking status was recorded for these two subgroups and was available in 97.2% (70/72) of carcinoid and in 80.0% (24/30) of LCNEC patients.

### 4.2 Immunohistochemistry

Immunohistochemistry (IHC) was performed using the automated immunostainer BenchMark Ultra and the OptiView detection system (Ventana Medical System Inc., Oro Valley, Arizona, USA). IHC for MTAP was performed retrospectively or during routine diagnostics using the mouse monoclonal anti-MTAP antibody clone 2G4, by Abnova (H00004507) at a dilution of 1:200. This antibody was chosen because of good experience in earlier studies^9,15^. Specimens with cytoplasmic expression of MTAP in viable tumor cells were labeled as “MTAP proficient”, irrespective of staining intensity. Viable tumor cells with complete lack of cytoplasmic MTAP expression in the presence of an internal positive control (any benign cell) were defined as “MTAP deficient”. In case of heterogeneous expression (unequivocal MTAP proficient and deficient tumor cells in the same tumor), the proportion of MTAP deficient tumor cells was given. Areas with mechanical artefacts or necrosis were not considered for evaluation. As recently described^15^, MTAP staining results are sensitive to insufficient formalin fixation in larger resection specimens. Therefore, MTAP was evaluated in better-fixed peripheral areas with special attention paid to internal positive controls (non-neoplastic cells within the tissue).

Selected carcinoids were additionally stained for OTP (orthopedia homeobox protein) and CD8 using the following antibodies and protocols: CD8 clone SP57, pre-diluted (Ventana, Tuscon, AZ) and OTP clone CL11225 (Sigma AMAb91696; dilution 1:50). Cases with nuclear expression of OTP in viable tumor cells were labeled as “OTP positive”; in heterogeneous cases, the percentage of OTP positive tumor cells was given. Tumor-infiltrating CD8 positive immune cells were quantitated by counting positive cells in an area of 2mm^2^ at 400x magnification (performed by LB). The area with the highest estimated density of CD8 positive cells was selected for this analysis. Data on the immunohistochemical expression of TTF-1 (Thyroid Transcription Factor-1), synaptophysin, and chromogranin A were collected retrospectively for the carcinoid samples.

### 4.3 Fluorescence in situ hybridization

In twelve MTAP deficient carcinoids with well-identifiable tumor areas on unstained empty sections, we performed fluorescence *in situ* hybridization (FISH) for evaluation of the 9p21 copy number status. We used a commercially available FISH assay (ZytoLight® SPEC CDKN2A/CEN 9 Dual Color Probe, Zytovision; #Z-2063) as previously described^8^. In the tumor cell nuclei, the complete absence of a 9p21 signal in the presence of a centromere 9 signal was considered ashomozygous deletion of 9p21. Adjacent nuclei of benign cells served as internal positive control.

### 4.4 Whole Exome Sequencing

#### DNA extraction from FFPE tissue blocks

FFPE tissue sections from eleven carcinoid specimens (six MTAP proficient, five MTAP deficient; Supplementary Table 1) and matched non-tumor samples were obtained from each tissue block and deparaffinized with xylene. The tumor areas were marked by a pathologist (LB) and dissected with a scalpel, and total DNA was extracted using the RecoverAll Total Nucleic Acid Isolation Kit for FFPE (Qiagen, 80234), following the manufacturer’s instructions. Final DNA concentrations were measured using the Qubit dsDNA HS Assay Kit (ThermoFisher, Q32851). A total of 500 ng of FFPE DNA from each sample was treated with uracil-DNA glycosylase (UDG) (NEB, M0280S) according to the manufacturer’s instructions. DNA samples were stored at –20°C prior to sequencing.

#### Whole exome sequencing and variant annotation

Whole exome sequencing was performed by CeGaT (Tübingen, Germany) using the Twist Human Core Exome + RefSeq + Mito-Panel kit (Twist Bioscience) for exome capture, according to the manufacturer’s instructions. Paired-end 100-bp reads were generated on an Illumina NovaSeq X Plus platform.

Raw FASTQ data processing workflow was adapted from *Roma et al.*^49^. Reads were aligned against the reference human genome GRCh38 using Burrows-Wheeler Aligner (BWA, v.0.7.12)^50^. SNVs and indels were called using MuTect2 (GATK v.4.1.4.1)^51^. To reduce false positive results from artifacts caused by formalin fixation, specific SNVs C:G>T:A with variant allelic fraction (VAF) less than 10% were discarded. We further excluded variants identified in at least two of a panel of 210 non-tumor samples^52^, including the non-tumor samples included in the current study. Variant annotation was performed by SnpEff software v.4.1^53^.

To ensure the genetic identity of tumor-normal pairs and exclude potential sample swaps or cross-contamination, we performed genotype verification using BAMixChecker^54^.

The heatmap of non-synonymous mutations was generated using the R package MAFTOOLS v.2.0.16^55^ by selecting the neuroendocrine lung cancer driver genes from *Alcala et al.*^18^.

#### Copy number alterations and clonality analysis

Allele-specific CNAs were identified using FACETS v.0.5.14^56^ and the log ratio relative to ploidy was used to call deletions, loss, gains and amplifications. Genes with total copy numbers greater than gene-level median ploidy were considered gains; greater than ploidy + 4, amplifications; less than ploidy, losses; and total copy number of 0, homozygous deletions. Cancer cell fraction (CCF) of each mutation was inferred using ABSOLUTE v1.0.6^57^. Solutions from ABSOLUTE were manually curated to ensure the solution matched the ploidy estimate generated by FACETS.

A mutation was classified as clonal if its probability of being clonal was > 50% or if the lower bound of the 95% confidence interval of its CCF was > 90%. Mutations were considered subclonal if they did not meet the mentioned criteria^58^.

### 4.5 Transcriptomic analysis

Five tumor specimens (two MTAP proficient (P1, P2) and three MTAP deficient (P8, P10, P12); Supplementary Table 1), for which fresh-frozen material was available, were used to perform bulk RNA sequencing. Initially, another MTAP proficient case was included (P3), which was subsequently excluded as it turned out to be a synovial sarcoma as supported by the detection of a SS18–SSX fusion. The tumor cell proportion of all five tested samples was above 80%. 120–200 µm OCT sections were obtained from each of the five samples. Isolation was performed using the AllPrep DNA/RNA Mini Kit (Qiagen, cat. no. 80204) following the manufacturer’s instructions. Briefly, tissues were homogenized in 350 µl RLT buffer using a stainless-steel bead in a TissueLyser (Qiagen, cat. no. 85210) for 5 min at a frequency of 18/s. The lysate was first passed through an AllPrep DNA spin column to bind genomic DNA, and the flow-through was collected for RNA purification. After addition of 70% ethanol, the sample was loaded onto an RNeasy Mini Spin Column and centrifuged at 13,000 rpm for 30 sec. Columns were treated with DNase I (RNase-Free DNase Set, Qiagen, cat. no. 79254) on-column to remove residual genomic DNA. After washing, RNA was eluted in 50 µl RNase-free water. RNA concentration was measured using the Qubit Fluorometer with the RNA High Sensitivity Assay Kit (Thermo Fisher Scientific, cat. no. Q32852). Samples were stored short-term at −20°C until shipment. RNA sequencing was performed by CeGaT GmbH (Tübingen, Germany) using 100 ng of total RNA per sample. Libraries were prepared using the KAPA RNA HyperPrep with RiboErase (HMR) kit with KAPA Globin Depletion Hybridization Oligos (F. Hoffmann-La Roche Ltd., Basel, Switzerland), according to the manufacturer’s instructions. Sequencing was carried out on an Illumina NovaSeq X Plus platform.

Reads were trimmed using Trimmomatic v.0.39^59^ and aligned to the GRCh38 human reference genome using STAR v.2.7.9.^60^. Transcript quantification was performed using RSEM v.1.1.3^61^.

A principal component analysis (PCA) was performed using the data normalized with the variance stabilizing transformation (VST) method as input and a plot was drawn using the ggplot2 R package v.3.5.2^62^.

Raw counts were normalized using the median of ratios method from the DESeq2 package v.1.42.1^63^ in R v.4.3. Differential expression analysis was performed using the DESeq2 Wald test and a volcano plot has been drawn using ggplot2 v.3.5.2^62^. Significant differential gene expression was considered for genes with a |LogFC| > 1.5 and adjusted *p* < 0.05.

Gene Set Enrichment Analysis (GSEA) was performed using the fgsea R package^64^. Genes were ranked according to a score calculated as sign(log2 Fold Change) × −log10(p-value). Enrichment was assessed against the Hallmark gene sets from MSigDB(v7.1)^65^. Only pathways containing between 15 and 500 genes were considered. Results were summarized using the Normalized Enrichment Score (NES) and Benjamini–Hochberg adjusted *p*-values and were visualized using ggplot2 v.3.5.2^62^.

The heatmap has been drawn using Pheatmap package v.1.0.13^66^ using the Z-normalized log-transformed gene expression values (Z-score) as input.

The tumor immune microenvironment on bulk RNAseq has been calculated using ConsensusTME software the TPM-normalized not log-transformed data using as input^67^. The scatter plots showing the correlation of MTAP with OTP expressions were drawn using the ggplot2 package v.3.5.2^62^ and Pearson’s correlation was calculated using the stat_cor() function implemented in the ggpubr package v.0.6.1^62^.

### 4.6 Deep Tissue Proteomics

#### Tissue sections on membrane slides

FFPE tissue sections (7 µm) from four MTAP proficient (P1, P2, P5, P7; Supplementary Table 1) and four MTAP deficient (P8, P10, P12, P13; Supplementary Table 1) carcinoid resection samples were mounted on Leica PPS frame slides (4 µm PPS membrane, steel frames; Leica, 11600294), and air-dried at room temperature. Sections were deparaffinized, rehydrated, stained with hematoxylin and eosin (H&E), and mounted using a xylene-based mounting medium with coverslips. Three reference markings per slide were lasered without perforating the membrane (10x HC PL Fluotar 10x/0.32 PH1 objective, laser power 15, Aperture 1, Speed 10, Head Current 100%, Pulse frequency 528) to facilitate accurate contour transfer between the digital annotation and laser microdissection microscope.

#### Slide scanning and processing

H&E-stained tissue sections were digitized using either ZEISS AxioScan at 20× magnification. After slide scanning, coverslips were removed by gentle agitation in xylene for 10 min at room temperature and the remaining mounting medium was carefully washed off. Slides were dipped in 100% ethanol, dried at 45 °C for 5 minutes, and stored at room temperature until laser microdissection.

#### Image analysis

Slide images were analyzed in QuPath (v.0.5.1, x64) and manually annotated by a board-certified pathologist (ND). For each tumor, annotations covered a cumulative area of 30,000 – 40,000 µm², corresponding to approximately 300 – 500 tumor cells. Annotated contours and reference markings were exported as GeoJSON files and converted into XML format compatible with LMD7 software using a custom Python script.

#### Laser microdissection

Tissue contours were excised and collected using the Leica LMD7 system with LMD software v.8.5.0.9136. Contours were dissected using a 20× objective (HC PL FLUOTAR 20×/0.40 CORR) in brightfield mode, with the following settings: power 36-40, aperture 1, speed 11, middle pulse count 2, final pulse 4, head current 100%, pulse frequency 555, and offset 119.

Contours were collected into twin.tec PCR plate 384 LoBind (Eppendorf, 0030129547) at a constant ambient temperature of 32 °C. Afterwards, collected contours were concentrated at the bottom of each well by adding 12 µL of 100% acetonitrile to rinse the walls, followed by vacuum drying at 60 °C for 20 min, 2000 rpm in a SpeedVac. All wells were microscopically inspected prior to sample preparation to ensure efficient tissue collection.

#### Sample preparation for MS

Samples in 384 well plates were incubated in 4 µL of 60 mM triethylammonium bicarbonate (TEAB), 0.1% DDM, 5 mM TCEP, and 20 mM CAA at 95 °C for 60 min in a PCR cycler. After addition of 1 µL 60% acetonitrile (final concentration 12% v/v), samples were incubated at 75 °C for another 60 min. Afterwards, proteins were digested with 4 ng/ul (each) trypsin and LysC at 37 °C overnight. Samples were then vacuum dried at 60 °C for 20 min at 2,000 g in a SpeedVac and resuspended in 20 µL of 2% acetonitrile (v/v), 0.1% trifluoroacetic acid (v/v), before completely loading onto C18 Evotips (Evotip Pure, EV2011, Evosep).

C18 Evotips loading was performed according to the manufacturer’s instructions. Briefly, 20 µL buffer B (99,9AC., 0.1% FA) was added to each C-18 tip and centrifuged at 800 g for 1 min. Subsequently, tips were activated in isopropanol for at least 20 s, and centrifuged again at 800 g for 1 min. Then, 20 µL buffer A (0.1% FA in H2O) was added and centrifuged at 800 g for 60 s. Digested samples were loaded onto Evotips, spun for 60 s at 800 g and washed once with 20 µL buffer A. Afterwards, loaded tips were topped with buffer A and stored at 4 °C until LC-MS/MS measurement was performed.

#### Liquid chromatography – mass spectrometry (LC-MS/MS) analysis

Peptide samples were analyzed by liquid chromatography-tandem mass spectrometry (LC–MS/MS) using an Orbitrap Astral mass spectrometer (Thermo Fisher Scientific). Peptides were separated using an EvoSep One liquid chromatography system equipped with an Aurora Elite 15×75 XT C18 column (IonOpticks). Samples were loaded using the standard EvoSep method and separated with a pre-designed 40 samples per day Whisper Zoom gradient at constant flow rate. Mobile phases consisted of solvent A (0.1% formic acid in water) and solvent B (0.1% formic acid in acetonitrile).

Data-independent acquisition (DIA) was performed in the Astral analyzer using sequential, non-overlapping isolation windows of 2 m/z across a precursor mass range of 380–980 m/z, with window placement optimization enabled. A total of 299 DIA scan events were collected per acquisition cycle. Fragment ions were generated by higher-energy collisional dissociation (HCD) using a normalized collision energy of 25% and detected over a scan range of 150-2000 m/z. MS2 scans were acquired with a scan time of 3 ms, a normalized automatic gain control (AGC) target of 300% (absolute target 30,000), and a single microscan. Multiplexed ion acquisition was disabled.

#### Data analysis of proteomic raw files

Raw spectral data were analyzed using Spectronaut (Biognosys, Schlieren; v.20.3.251215.x). Spectra were searched against the human Swiss-Prot FASTA database (exported Nov. 2021, 20,360 entries) in library-free directDIA mode. Carbamidomethylation of cysteins was set as fixed modification, n-terminal acetylation and methionine oxidation were set as variable modifications. A minimum peptide length of seven amino acids was required. Enzyme specificity was set to trypsin and LysC with up to two missed cleavages. The final library comprised 167,826 precursors, 143,309 peptides, and 9223 protein groups (protein LFQ in automated setting).

#### Quantification and statistical analysis

Proteomics data were analyzed using Perseus v.2.1.3.0^68^ and R v.4.2.2. Identified proteins were filtered to retain proteins with at least 70% non-missing values in at least one group (MTAP proficient and deficient), and intensities were median aggregated across technical replicates, filtered for detection in at least 2 of 3 replicates.

Differential expression analysis was performed using the limma R/Bioconductor package^69^. A linear model was fitted to the data for each protein using the limma lmFit function, with an experimental design matrix specifying the comparison between MTAP-proficient and MTAP-deficient samples. Model coefficients were estimated and variance was moderated using empirical Bayes statistics (eBayes). Differential protein abundance was determined using thresholds of an absolute log2 Fold Change (|log2FC|) > 1 and a Benjamini–Hochberg adjusted *p*-value < 0.05 to control the false discovery rate (FDR).

#### Gene Set Enrichment Analysis

Gene Set Enrichment Analysis (GSEA) was performed using the fgsea R package^64^. Genes were ranked according to a score calculated as sign(log2 Fold Change) × −log10(p-value). Enrichment was assessed against the Hallmark gene sets from MSigDB(v7.1)^65^. Enrichment results were summarized using the normalized enrichment score (NES) and Benjamini–Hochberg adjusted p-values to control the false discovery rate (FDR), and were visualized using ggplot2 v.3.5.2^62^.

### 4.7 Methylation analysis

DNA methylation profiling was performed on the same five RNAseq samples data (n=2 MTAP proficient, n=3 MTAP deficient) for which frozen material was available. Genomic DNA was processed at Life&Brain GmbH (Bonn, Germany) using the Illumina Infinium MethylationEPIC v2 array, covering 937,690 CpG sites (according to the manufacturer’s instructions).

Bioinformatic analysis was conducted in an R environment v.4.3. Raw intensity data (*IDAT* files) were processed and normalized using the SeSAMe package v.1.20.0^70^ with the hg38 (GRCh38) assembly as reference genome. To ensure high data quality, beta values were subjected to a rigorous filtering pipeline: CpG sites were retained if they had a missingness rate ≤20%, while probes with negligible variance or extreme beta values (0 or 1) in more than 10% of samples were excluded. Following these filtering steps, 862,719 CpG probes were retained for downstream analysis. Global data structure and sample relationships were assessed using Principal Component Analysis (PCA) and Pearson’s correlation matrices. Differential methylation analysis between MTAP-proficient and MTAP-deficient groups was performed by converting beta values to M-values and applying linear models via the limma package v.3.58.1^69^ with empirical Bayes smoothing (*eBayes*). Finally, a targeted analysis was conducted on a predefined set of CpG probes of interest. These probes were selected based on their high variability across multiple cancer types near MTAP and OTP promoter regions in the TCGA Pan-Cancer Atlas (Xena, GDC-PANCAN dataset), as well as previously validated probes from published literature ^30^. Methylation levels for these sites were visualized and compared using statistical boxplots generated with ggplot2 v.3.5.2^62^. The heatmap showing all the CpG sites within the *MTAP* gene was generated using Pheatmap R package v.1.0.13^66^.

### 4.8 Publicly available data

Raw count data from a previously published study on neuroendocrine lung cancer by *Gabriel et al.*^20^ were downloaded from the DRMetrics GitHub repository (https://github.com/IARCbioinfo/DRMetrics). The dataset includes gene expression profiles from 75 typical carcinoids, 37 atypical carcinoids, 69 LCNEC, 51, SCLC, and 3 supra-carcinoids ^18,20^. Raw counts were normalized using the median-of-ratios method implemented in the DESeq2 R package v.1.42.1^63^. Heatmaps were generated using the Pheatmap R package v.1.0.13^66^, based on Z-score normalized log-transformed gene expression values. Additional independent transcriptomic datasets were retrieved from public repositories. SCLC transcriptomic data^46^ were obtained from the Gene Expression Omnibus (GEO) under accession number GSE60052 in log-normalized format. Carcinoid transcriptomic data were retrieved from GEO under accession number GSE118131^17^ in log2TPM normalized format. Raw transcriptomic data for LCNEC were downloaded from the supplementary material of *George et al*.^43^ and normalized using the median-of-ratios method implemented in DESeq2 v.1.42.1^63^, followed by log transformation. For comparative expression analyses across datasets, boxplots were generated using ggplot2 v.3.5.2^62^, and statistical significance was assessed using the t-test implemented in the ggsignif R package v.0.6.4^71^. DNA methylation data in *Beta* value format from a previously published study^18^ were downloaded from the European Phenome-Genome Archive (EGA) under accession number EGAD00010001720. Additional DNA methylation data on carcinoids in *Beta* value format were downloaded from GEO under accession number GSE118132.

## Abbreviations

NEN: pulmonary neuroendocrine neoplasms
IHC: immunohistochemistry
FISH: fluorescence in situ hybridization
WES: whole exome sequencing
LCNEC: large cell neuroendocrine carcinomas
SCLC: small cell lung cancers
ICI: immune checkpoint inhibitors
NSCLC: non-small cell lung cancer
MTAP: S-methyl-5’-thioadenosine phosphorylase
MTA: methylthioadenosine
CNV: copy number variations
TMB: tumor mutational burden
OTP: orthopedia homeobox protein
PCA: principal component analysis
RNA-seq: RNA sequencing
lncRNAs: long non-coding RNAs
miRNAs: microRNAs
aSCLC: atypical SCLC

## Data Availability

The WES and bulk RNAseq data generated during this study have been submitted to the European Genome-Phenome Archive (EGA) under the accession numbers EGAS50000001683 (bulk RNAseq), EGAS50000001684 (WES). Proteomics data generated during this study have been submitted to the EBI PRIDE repository under accession number PXD074011. DNA methylation data generated during this study will be submitted to the NCBI Gene Expression Omnibus (GEO) repository. The datasets generated and analyzed during the current study will be publicly available upon publication.

## Acknowledgments

We acknowledge CeGaT (Tübingen, Germany) for its technical expertise in sequencing; Niklas Simon for his assistance in cutting sections from frozen tumor tissues. The Data Access Committee of the Medical Faculty of the University of Basel (MF-DAC) for their support in providing the authorization to submit the WES and RNAseq data to the European Genome-Phenome Archive; Calculations were performed at sciCORE (http://scicore.unibas.ch/) scientific computing center at the University of Basel.

Magdalena M. Brune is supported by the Swiss National Science Foundation (Grant Number <u>P500PM_217648</u>).

Nikolaus Deigendesch is funded by the Nuovo-Soldati Foundation

Jürgen Hench is funded by the Gertrude-von-Meissner Stiftung, Basel, Switzerland.

## Author contributions

M.M.B, L.R., and L.B. conceived the study. L.B. supervised the study. L.R., N.D., F.M., S.U., P.H., J.P., J.H. and I.B.H. designed and conducted experiments. M.M.B., L.R., S.U., N.D., and L.B. analyzed data and performed data visualization. M.M.B., K.M.K., D.K., C.P. S.R.O., S.S.P., M.H. and L.B. contributed to patient enrollment, collection of samples and clinical data, and provided clinical advices. M.M.B., O.C., J.K., K.M.K, S.S.P., M.H. and L.B. performed the histopathological evaluation. M.M.B., L.R. and L.B. wrote the initial manuscript and all the authors contributed to the final version. All authors have seen and approved the manuscript.

