## Supplementary Material for "MTAP deficiency is a novel biomarker in neuroendocrine neoplasms of the lung"

**MTAP deficiency is a novel biomarker in neuroendocrine neoplasms of the lung**
**Supplementary Material List**

Supplementary Figure 1: Transcriptomic profiling of neuroendocrine lung tumors.

Supplementary Figure 2: Genomic and epigenomic features of MTAP proficient and MTAP deficient carcinoids.

Supplementary Figure 3: Transcriptomic landscape of MTAP proficient and MTAP deficient carcinoids.

**Supplementary Tables:**

Supplementary Table 1: Metadata of carcinoid samples

Supplementary Table 2: List of mutations identified by WES

Supplementary Table 3: WES quality metrics

Supplementary Table 4: Tumor purity and ploidy estimated from WES

Supplementary Table 5: Differentially expressed genes in MTAP-proficient versus MTAP-deficient carcinoids

Supplementary Table 6: Differentially expressed proteins in MTAP-proficient versus MTAP-deficient carcinoids

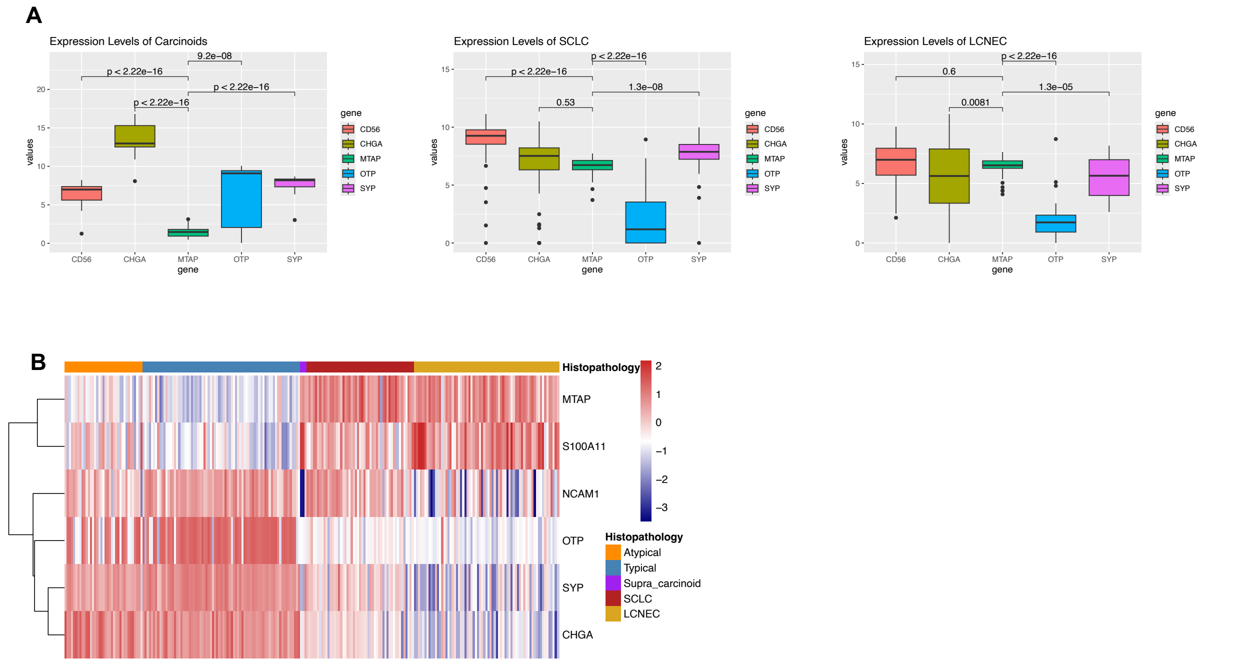


**Supplementary Figure 1. Transcriptomic profiling of neuroendocrine lung tumors.**

**A)** RNA Expression of *CD56*, *CHGA*, *MTAP*, *OTP*, and *SYP* in lung tumors. Left: carcinoids (n=30) from *Laddha et al*[1]; center: small cell lung carcinomas (SCLC, n=79) from *Jiang et al*[2].; right: large cell neuroendocrine carcinomas (LCNEC, n=66) from *George et al*. Data are log-transformed.

**B)** Heatmap of transcriptomic data from Gabriel *et al*[3]. dataset. The x-axis shows five supervised clusters: Typical Carcinoids (n=75), Atypical Carcinoids (n=37), Supra Carcinoids (n=3), SCLC (n=51), and LCNEC(n=69). Gene expression data are Z-normalized and log-transformed.

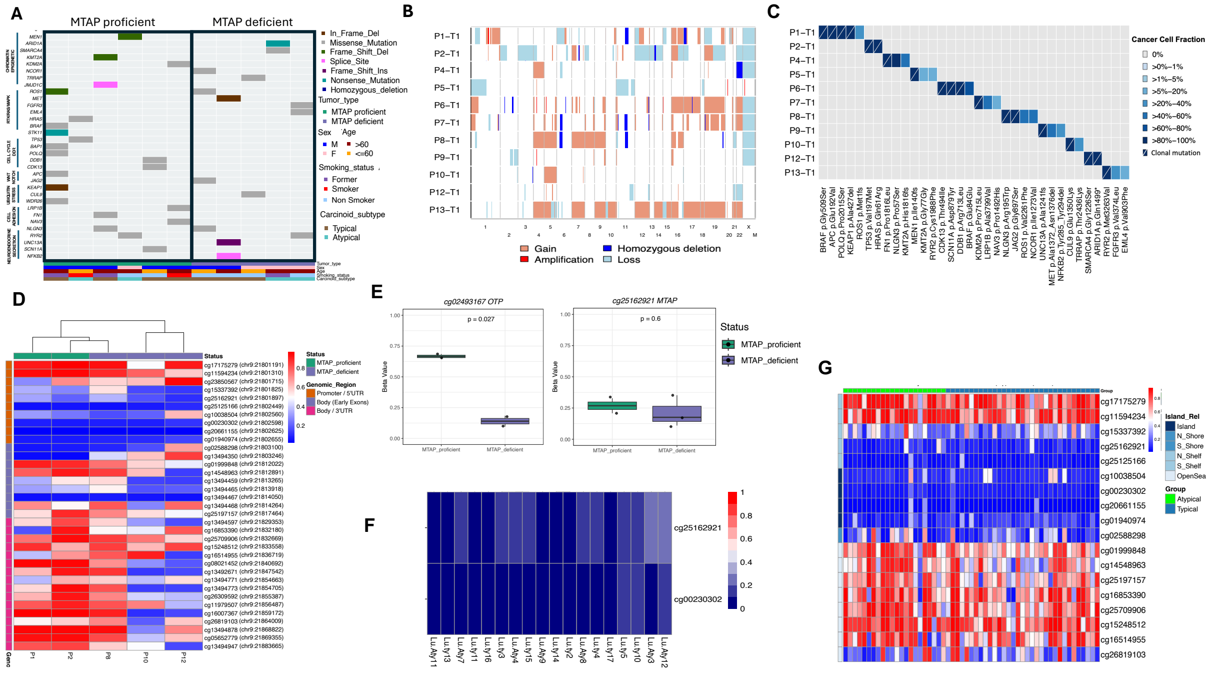


**Supplementary Figure 2: Genomic and epigenomic features of MTAP proficient and MTAP deficient carcinoids.**

**A**) Non-synonymous mutations plot of microdissected carcinoids from eleven patients. Cancer driver genes were defined according to IntOGen and Bailey *et al.*[4], and carcinoid driver genes according to Alcala *et al*[5]. The black frame separates MTAP proficient from MTAP deficient samples.

DDR = DNA Damage Response.

**B)** Heatmap illustrating the copy number variation (CNV) profile of all samples.

**C)** Heatmap showing the cancer cell fraction (CCF) of non-synonymous driver mutations. Clonal mutations are indicated by a diagonal line; amino acid changes are shown close to the genes.

**D**) Heatmap showing *MTAP* methylation status (n=5) using Illumina EPIC microarray V2 data performed on our cohort. Beta values <0.2 (blue) indicate hypomethylation and >0.6 (red) indicate hypermethylation.

**E)** Boxplots showing the methylation status of CpG probes associated with *MTAP* (left, cg25162921) and *OTP* (right, cg02493167) promoters. Methylation levels are expressed as beta values, ranging from 0 (unmethylated) to 1 (fully methylated). Individual data points represent independent samples, categorized by MTAP status.

**F)** Heatmap showing *MTAP* promoter CpGs (cg00230302 and cg25162921) methylation analysis in 18 carcinoid samples from publicly available data. Beta values <0.2 indicate hypomethylation and >0.6 indicate hypermethylation.

**G)** Heatmap showing *MTAP* methylation status (n=55) using Illumina EPIC microarray (850K) data from Alcala *et al.*[5] Beta values <0.2 (blue) indicate hypomethylation and >0.6 (red) indicate hypermethylation.

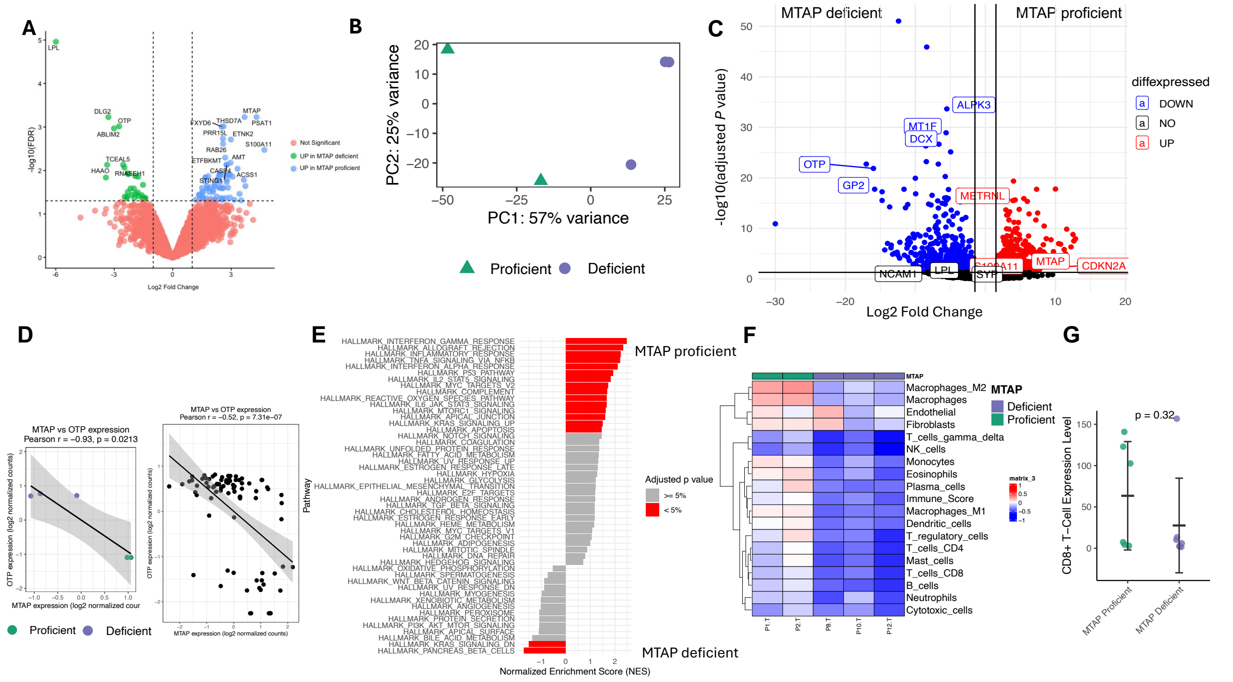


**Supplementary Figure 3: Transcriptomic landscape of MTAP proficient and MTAP deficient carcinoids.**

**A**) Volcano plot of differentially expressed proteins in carcinoids comparing MTAP proficient (n=4, right) versus MTAP deficient samples (n=4, left) by proteomics. Significantly up-regulated proteins (adjusted *p* < 0.05) are shown in blue, down-regulated in green, and non-significant in pink.

**B)** Principal component analysis (PCA) showing transcriptomic profiles based on the 100 most variable genes.

**C)** Volcano plot of differentially expressed genes in carcinoids comparing MTAP proficient (n=2, right) versus MTAP deficient samples (n=3, left) by transcriptomics. Significantly up-regulated genes (adjusted *p* < 0.05) are shown in red, down-regulated in blue, and non-significant genes in black.

**D)** Scatterplots showing *Pearson's* correlation between *MTAP* and *OTP* expression in transcriptomics (top left, n=5), and transcriptomic data from *Gabriel et al*[3]. (right, n=82). *Pearson's* correlation coefficient *(r)* and *p* are indicated on each plot.

**E)** Gene set enrichment analysis (GSEA) showing hallmark pathways in MTAP proficient (top) versus MTAP deficient (bottom) samples based on transcriptomics. Significantly deregulated pathways (adjusted *p* < 0.05) are highlighted in red.

**F)** Heatmap showing the tumor microenvironment (TME) score comparing MTAP proficient carcinoids (n=2) versus MTAP deficient carcinoids (n=3).

**G)** Jittered strip plot displaying the distribution of CD8+ T-cell expression metrics between MTAP-proficient (n=6) and MTAP-deficient (n-7) cohorts. Individual data points represent independent biological samples. Central horizontal lines and error bars indicate the mean ± standard deviation (SD) for each group. Statistical significance was evaluated using a two-tailed t-test.
